## Supplementary Information for "LLPS REDIFINE allows the biophysical characterization of multicomponent condensates without tags or labels"

### 1. NMR diffusion experiments with complex biomolecular systems

In the traditional NMR diffusion experiment, a signal of an analyte is mapped as a function of the gradient amplitude applied in the pulsed gradient spin echo (PGSE) experiment.[1,2] The use of gradients can spatially label molecules depending on their position in the NMR tube. Their translational diffusion will lead to a signal attenuation that depends on experimental parameters and diffusion rates (Methods, Supplementary Fig. 1a) and it follows a simple mono-exponential decay (Supplementary Fig. 1b, Methods - Eq. 1) providing a quantitative measure of the displacement. Often, this decay is visualized on a semi-logarithmic plot (Supplementary Fig. 1b, right), where a slope of the line simply represents the diffusion coefficient (Methods - Eq. 3). Note that due to technical limitations of the applied gradient strength, very slow diffusion rates such as those associated with molecular complexes and biomolecules in a viscous environment ($D < 5\times10^{-12}$ m$^2$/s) cannot be measured on conventional NMR probes and appear as flat non-decaying curves (Supplementary Fig. 1b).

Generally, proteins and large RNAs inside the viscous droplets undergo very slow diffusion. However, when multiple exchanging species are present in the sample (or molecules coexisting in multiple states) with partially or fully overlapping NMR resonances, the PGSE curve exhibits a complex decay affected by both species (Supplementary Fig. 1c). The chemical exchange imparts the information about the apparently invisible diffusion coefficient $D_2$ onto

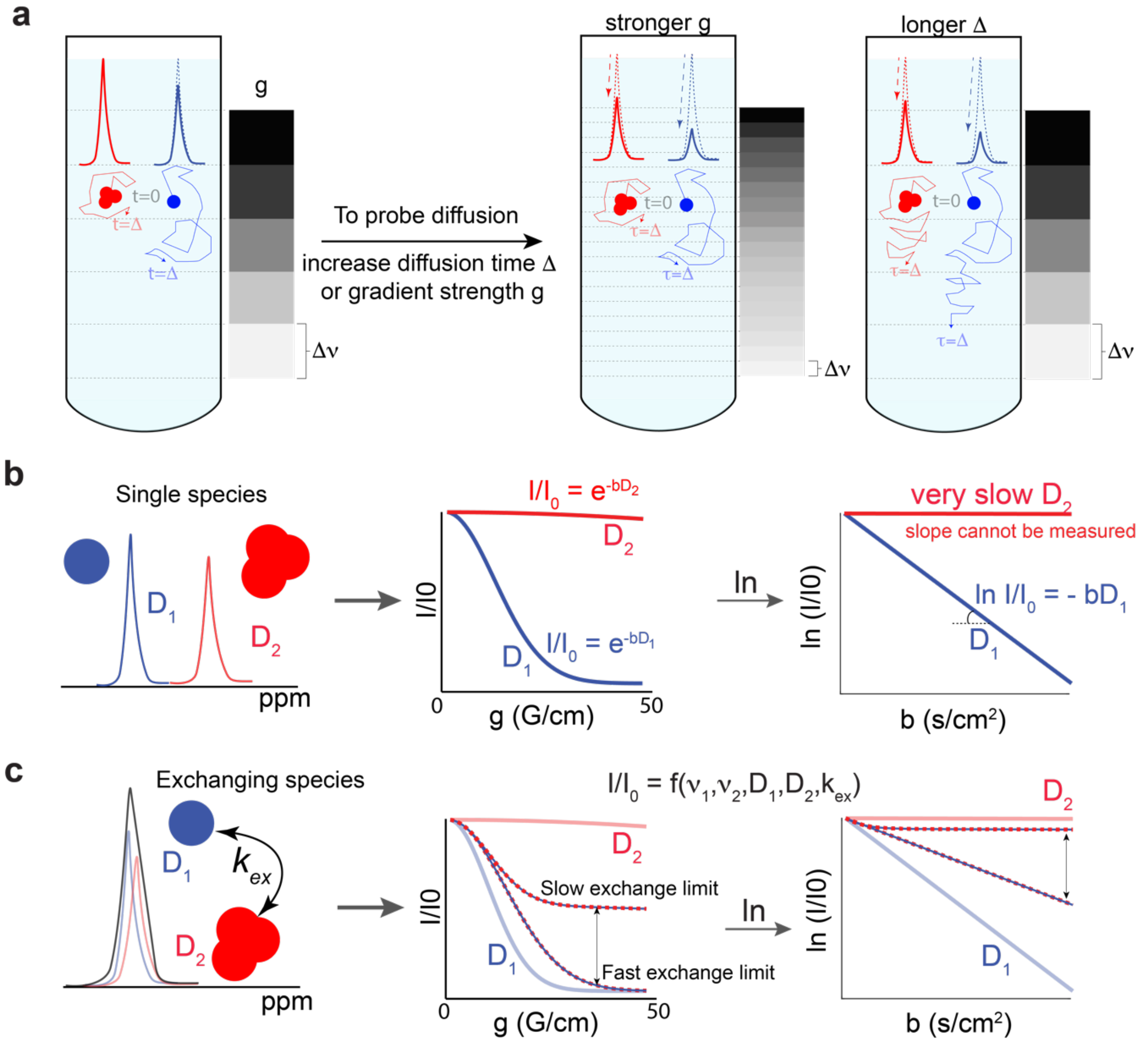

**Supplementary Fig. 1: Principles of diffusion NMR experiments and the effect of exchange in complex systems. a** Application of a gradient along *z*-axis can effectively "divide" an NMR sample in slices along the same axis. Translational diffusion of a molecule to other neighboring slices will manifest as the signal decay. To quantify diffusion coefficient, traditionally one need to acquire multiple spectra for increasing gradient strength (which makes thinner slices) until all molecules left their initial slice. Note that also increasing diffusion time can yield the same effects since a molecule is allowed longer time to diffuse. **b** Simulated attenuation of the PGSE signal due to a diffusion of single species in solution decays mono-exponentially and appears as a line on semi-logarithmic plot. Very slow diffusion coefficients cannot be probed at most conventional instruments and appear flat in PGSE experiments. **c** The presence of multiple species exchanging with each other encodes an exchange-averaged diffusion information on the NMR diffusion decay. The observed diffusion curve will strongly depend on the exchange rate. Note the large deviation of the resulting diffusion curve from both dilute and condensed phase diffusion. Diffusion coefficients are randomly chosen to be $D_1 = 9\times10^{-11}$ m$^2$/s and $D_2 = 3\times10^{-13}$ m$^2$/s, illustrating the typical 2 orders of magnitude diffusivity difference in biomolecular condensates.

the observed diffusion curve, resulting in a trajectory appearing between the slow and fast exchange limits. On the semi-logarithmic plot (Supplementary Fig. 1c, right) this exchange-averaged diffusion decay (defined with an apparent diffusion coefficient) deviates from the diffusion of both dilute and condensed populations not allowing diffusion constants to be determined.

### 2. The framework of LLPS REDIFINE

The foundation of LLPS REDIFINE is illustrated by the model in Supplementary Fig. 2a. The model of chemical exchange equilibrium between different-diffusing species shown in Fig. 1c can be incorporated within the framework of the McConnell equations[3,4] (Methods). Due to restricted diffusion, we defined an apparent diffusion coefficient in a condensed phase that depends on droplet size and the choice of diffusion time (Methods). The apparent diffusion constants and their effect on diffusion in the condensed phase are shown in Supplementary Fig. 2b. For small droplet radiuses and long diffusion times, the apparent diffusion coefficient can be almost 3 orders of magnitude slower than the intrinsic diffusion coefficient while approaching its values for large radiuses emulating almost continuous condensed phase. The influence of permeability and resulting chemical exchange on the PGSE curves of molecules in condensed and dilute phases is presented in Supplementary Fig. 2c. Although the slow diffusion of protein in the condensed phase cannot be accurately measured by NMR PGSE experiment on conventional probes, the chemical exchange effectively mixes the information about the restricted diffusion with the NMR PGSE visible diffusion of the molecules in dilute phase. This leads to apparently faster decays allowing us to extract the information about condensed phase. What is more, NMR diffusion curves are strongly affected by the population of the protein in the condensed phase (Supplementary Fig. 2d). Using our model, we could simulate the behavior of the FUS NTD protein data shown in Fig. 1b to the finest details. Supplementary Fig. 2e shows the importance of including droplet size and permeability. Note the difference between considering two diffusion coefficients in slow exchange (typical for FUS) and the LLPS REDIFINE model. Using the former, diffusion coefficients of molecules in condensed and dilute phases and their populations could not even be approximated. Our simulations show that the PGSE curves are also significantly influenced by the choice of diffusion encoding time. From simulation results, we concluded that multiple diffusion decay curves spanning from short (30-50 ms) to long diffusion times (700-1000 ms), each carrying unique information about the chemical exchange and restricted diffusion (Supplementary Fig.

2f), are required to unambiguously describe all the parameters of biphasic condensates presented in the model. It is this rich set of diffusion information that allows LLPS REDIFINE to quantitively interpret the experimental data acquired on biomolecular condensates.

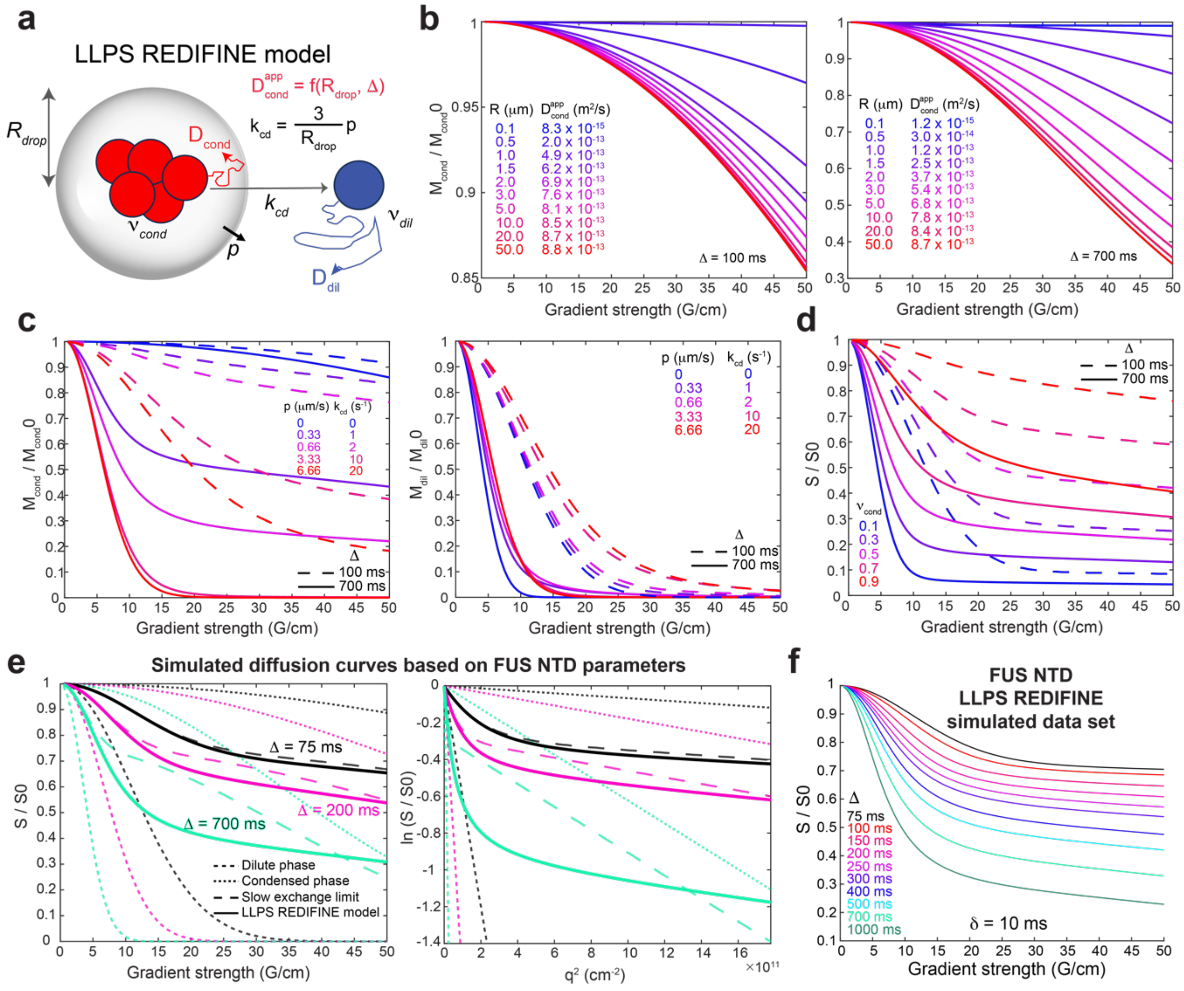


**Supplementary Fig. 2: Translating the restricted diffusion model with exchange into LLPS REDIFINE – simulations. a** The illustration of the REDIFINE model considering the effects of multiple parameters including corresponding phases' diffusion coefficients and populations, permeability and droplet size on the diffusion behavior in condensates. **b** Influence of droplet size on apparent diffusion in condensed phase simulated for two diffusion times, 100 and 700 ms. To isolate only the effect of droplet size, permeability is put close to zero (0.000333 µm/s). **c** Effect of permeability and resulting chemical exchange on the diffusion of molecules in condensed and dilute phase. Condensed phase decay becomes apparently faster with increasing exchange while opposite is true for the dilute phase. This is effective mixing of the information between phases. $R_{drop}$ is chosen to be 1 µm. **d** Influence of populations on the observed diffusion curves. $R_{drop}$ and p were 1 µm and 0.33 µm/s respectively. **e** Reproducing the diffusion curves observed for FUS NTD in Fig. 1b. Note that individual diffusion coefficients and populations cannot be approximated from the diffusion data without implementing LLPS REDIFINE model. **f** Full REDIFINE data sets including multiple diffusion curves acquired for different diffusion times. The data set with at least 6 curves spanning from short (50-75 ms) to long diffusion times (500-1000 ms) is needed to uniquely define the 5 parameters of the system ($D_{dil}$, $D_{cond}$, $\nu_{drop}$, $R_{drop}$, and p). The diffusion coefficients used in this simulations are chosen to match the ones of FUS NTD, $D_{dil} = 8\times10^{-11}$ and $D_{cond} = 8.9\times10^{-13}$.

### 3. LLPS REDIFINE – experimental considerations

To acquire the experimental data sets, we used a pseudo-3D stimulated echo NMR experiment with bipolar gradients (PGSTE) where both the gradient strength and the diffusion delay are incremented as shown in Supplementary Fig. 3a. The theory shown in methods section for PGSE experiments apply here as well. Stimulated spin-echo probes diffusion while magnetization is stored on the *z*-axis, allowing prolonged diffusion encoding times as it relies on long longitudinal relaxation. Therefore, LLPS REDIFINE provides also a $T_1$ weighting, however as longitudinal relaxation of observable non-labile protons from flexible regions in condensed and dilute phase is nearly identical (Supplementary Fig. 3b,c), relaxation can be disregarded from the model. This is done by normalizing the data sets for different diffusion times to unity.

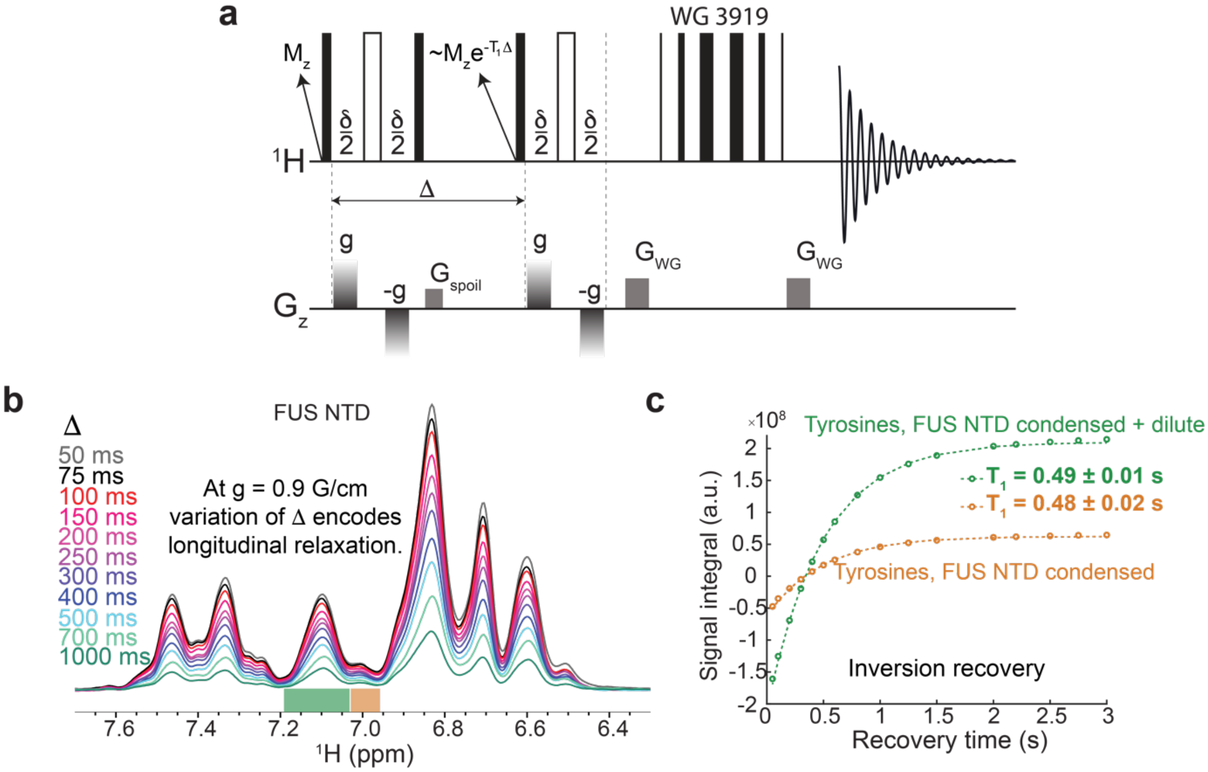


**Supplementary Fig. 3: Stimulated spin echo with bipolar gradients is used to acquire LLPS REDIFINE data. a** Stimulated gradient spin echo (PGSTE) is a preferred experiment to acquire NMR diffusion curves because it relies on slower $T_1$ relaxation allowing us to probe longer diffusion encoding time. **b** Signal decay of proton signals experienced by biphasic FUS NTD as a result of varying diffusion time has a $T_1$-weighting. Resonance highlighted in green represents tyrosines overlapping signals from protein in dilute and condensed phase in this FUS NTD biphasic sample. Resonance highlighted in orange contains signal solely from condensed protein in the droplets stuck to the wall of NMR tube (assigned based on systematic chemical shift from the main resonance).[16] **c** Independent inversion recovery experimental results shows that the longitudinal relaxation constants of non-exchangeable protons are the same for the protein in condensed and dilute phases based on excellent monoexponential fitting and identical relaxation values.

### 4. Diffusional kurtosis effect

The presence of chemical exchange between dilute and condensed phase leads to complex diffusion decays. On the semi-logarithmic plot (Supplementary Fig. 4a) the exchange-averaged diffusion decay (defined with an apparent diffusion coefficient) manifests as a deviation from the linear trend, often described as a kurtosis effect[5]. Kurtosis, denoted by the dimensionless parameter K, is a statistical metric[5] for quantifying the shape of a probability distribution. By definition, a Gaussian distribution has K equal to 0. Distributions that are more "peaked"

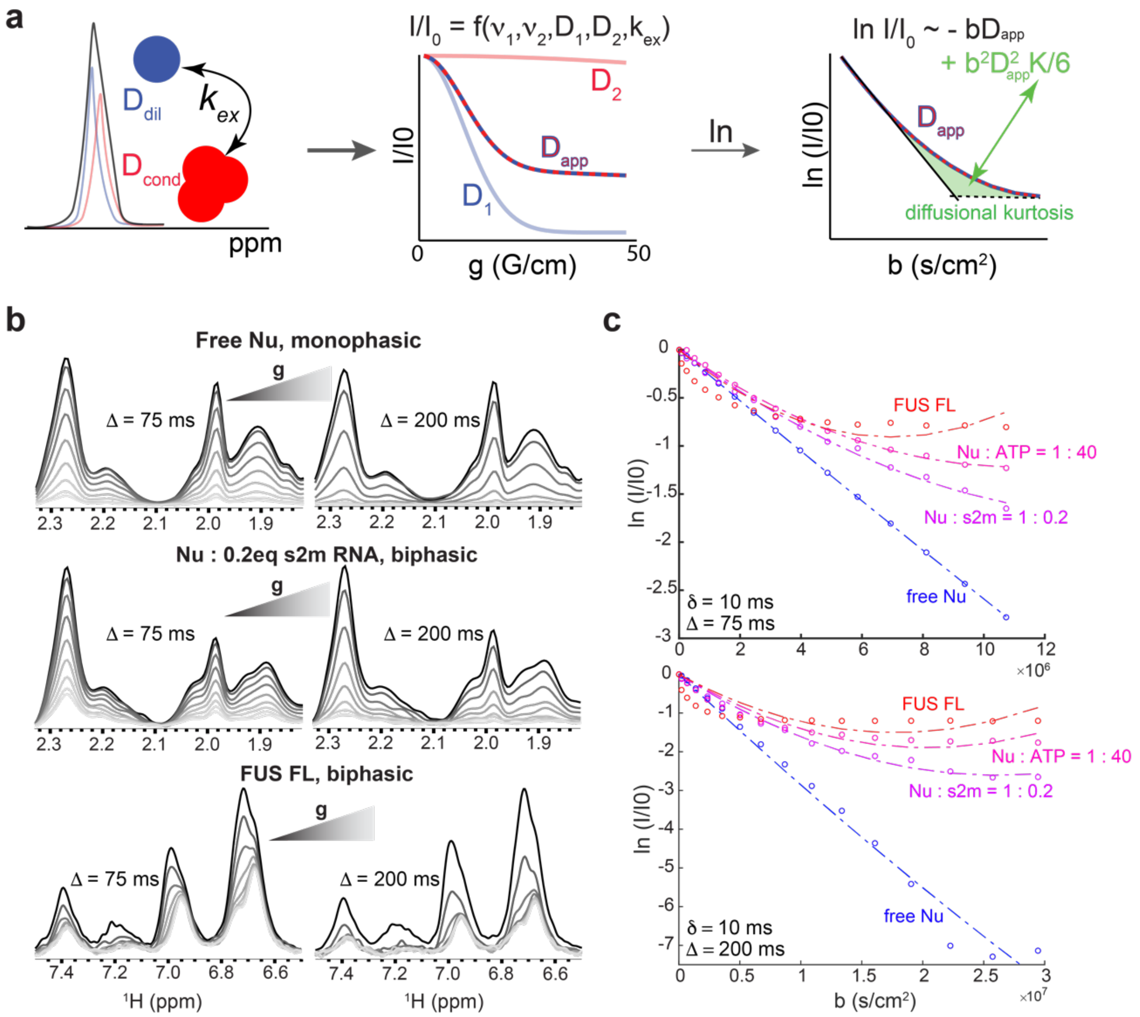


**Supplementary Fig. 4: Biphasic condensates undergo non-gaussian diffusion described with the kurtosis effect. a** Chemical exchange between dilute and condensed phase encodes an exchange-averaged diffusion information on the NMR diffusion decay. The observed diffusion curve will strongly depend on the exchange rate (droplet size and permeability). Note the large deviation from the linearity when the curve is shown as a $\ln\frac{I}{I_0}$ dependence of the diffusion factor b, described by kurtosis effect (Methods). **b** NMR signal decays of dispersed Nu, and biphasic Nu : RNA and FUS samples with respect to increasing gradient strength in PGSE experiment. Note the different decay trends when diffusion time Δ is increased from 75 ms to 200 ms. **c** All the condensate samples show significant diffusional kurtosis effect (deviation from linearity) compared to the free Nucleocapsid sample that experience pure mono-exponential decay. The kurtosis effect varied from 0.06 for free Nu to 1.66 for FUS condensate at 75 ms diffusion time. It is noteworthy that even Nu condensates with different nucleic acids show distinct kurtosis values (K = 0.87 for Nu:s2m and K = 1.23 with Nu:ATP). Both diffusion coefficients and kurtosis factors were notably different when using 200 ms diffusion time.

typically have a positive kurtosis (K > 0). Positive kurtosis reflects various diffusion environments experienced by diffusing molecules as they encounter barriers, [6] move between compartments, and undergo chemical exchange.[7,8] [9,10] It can also be a result of intermolecular interactions present in the system. A molecule diffusing with positive kurtosis would typically not travel as far in a given time interval as one that followed Gaussian statistics. In case of diffusional kurtosis, as it is shown in Supplementary Fig. 4a, the PGSE decay deviates from linearity on a semi-logarithmic plot. A traditional way to account for kurtosis is to perform a one-step Taylor series expansion of the exponent in the standard diffusion signal equation (Eq. 1), yielding

$$\frac{I}{I_0} = e^{-b \cdot D_{app} + b^2 D_{app}{}^2 K/6} \tag{S1}$$

As in the case of diffusional kurtosis, diffusion coefficient becomes a function of experimental parameters, obstacle size or interaction affinity, apparent diffusion coefficient $D_{app}$ must be introduced.[11–15]

It has been previously shown in MRI that diffusional kurtosis of water is very sensitive to the tissue type and even could be used to assess structure abnormalities in brain associated to neuropathologies.[7,8] Supplementary Fig. 4b shows the behavior of the NMR signals of different condensates (stabilized in agarose hydrogel)[16] subjected to diffusion NMR experiments. While signals stemming from free Nu in dispersed sample decay to zero at high gradient strengths, the peaks detected in the biphasic samples of Nu (in complex with RNA) and FUS (free) remain detectable even at high gradient values. For FUS protein, this signal plateau was correlated in our previous publication with the fraction of slowly diffusing protein in the condensed phase.[16] Similar behavior was initially anticipated for the biphasic Nu sample, however we noticed that the Nu condensate signals were experiencing a much steeper diffusion decay without an obvious plateau. As multi-component systems (protein, RNA and two phases), all the condensed samples show a significant kurtosis effect that intriguingly depends on the choice of diffusion time (Supplementary Fig. 4c). This implies the presence of restricted diffusion for the biomolecules and chemical exchange between the phases.

### 5. Scope of LLPS REDIFINE

LLPS REDIFINE method proved applicable for the most samples examined in this study. This includes almost 15 different samples prepared using 5 different proteins having a wide range of biophysical properties (Extended Data Table 1). Due to very fast interphase exchange, LLPS REDIFINE only failed to determine droplet parameters for PTBP1:3xUCUCU condensate samples. Chemical exchange of more than 30 $s^{-1}$ appears fast on the diffusion time scale averaging out the effects of restricted diffusion in the condensed phase and preventing unambiguous fitting of the LLPS REDIFINE datasets. Concluding from the simulation and experimental data, ideal chemical exchange for LLPS REDIFINE should fall between 0.5 $s^{-1}$ and 30 $s^{-1}$. While the equal population of the species in the two phases gives LLPS REDIFINE the best sensitivity to distinguish accurately condensate parameters, this is often not the case if one of the phases is populated less than 10 %. Then not all parameters can be unambiguously determined. However, the performance of LLPS REDIFINE will also depend on the signal-to-noise ratio (SNR) of signals detected in NMR. This is why very good SNR ratio when detecting water allowed LLPS REDIFINE to determine parameters with good accuracy based on only 3% water in the condensed phase.

To avoid the interference from chemical exchange that labile protons in protein undergo with bulk water,[9,10] only non-exchanging NMR peaks should be integrated in LLPS REDIFINE. Throughout this study, we usually integrated aromatic protons (FUS NTD, full-length FUS, A6 RNA) or aliphatic protons (DDX4, Nucleocapsid, PTBP1).

It appeared that the two-pool model is sufficient to explain most of the data that we acquired. For protein-RNA condensates, we tried to extend the number of species in the model, for example, to take into account protein in free and bound state in both phases. It is generally straightforward to implement this in Eq. 7, however, with more pools corresponding system of coupled differential equations cannot be solved analytically and has to be fitted numerically which increases the calculation time and complexity of the multiparametric fitting procedure. As this hasn't improved the fitting of the data acquired on Nu:A6 condensates (Fig. 3b) at shorter diffusion times we decided to fit all the protein-RNA condensates data with the two-pool approximation (one species in dilute and one in condensed phase).

With structured protein:RNA condensates, another issue can be the broadening of the signal upon binding. While the peaks originating from the folded domains that directly bind to RNA will be most likely broadened beyond detection, signals from flexible, disordered regions remain NMR observable (Supplementary Fig. 5a). As illustrated for Nucleocapsid construct 247-419 consisting of a structured RNA binding domain and intrinsically disordered tail, signals from IDR have very similar linewidths in dilute and condensed phase (Supplementary Fig. 5b). This is also the case for full-length Nucleocapsid, PTBP1 and FUS samples. Undetectable NMR signal of protein or RNA upon condensate formation (as for s2m RNA upon binding to Nucleocapsid protein) limits the applicability of LLPS REDIFINE.

Regarding soluble complexes, the methodology is most sensitive when diffusion coefficients of the exchanging species are as different as possible. Furthermore, as in the case of condensates, the chemical exchange ($k_{off}$ rate) needs to be in an intermediate regime on the diffusion time scale (residence time 30-1000 ms). This is often the case with RNA binding proteins and their complexes with RNA.

Even the presence of two non-exchanging species could lead to a kurtosis effect which was the case with a 2-week-old Nucleocapsid sample containing degradation proteoforms (Supplementary Fig. 6a,b). Having overlapping NMR signals and showing diffusional kurtosis served as a negative control for our method. REDIFINE methodology resolved that the species don't exchange with each other and determined the population and diffusion coefficients of the individual components. Even at three different temperatures altering diffusion behavior and kinetics, REDIFINE detected no chemical exchange between two species and kept the populations constant (Supplementary Fig. 6c). The only parameters that changed with temperature were diffusion coefficients which was an expected behavior.

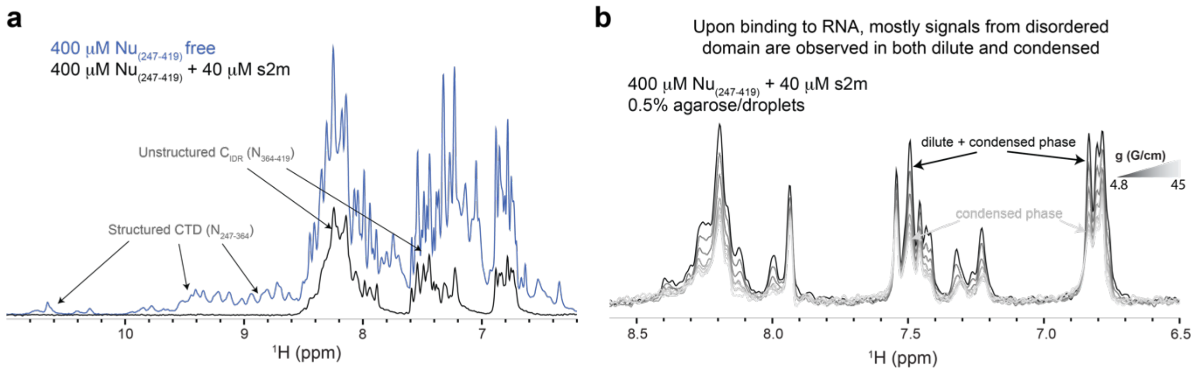


**Supplementary Fig. 5: IDRs remain NMR observable in the condensed phase. a** Even though signal of structured domains can broaden beyond detection because of slower tumbling upon protein binding and/or increased viscosity in condensed phase, IDR regions remain flexible. 1D spectra show the comparison of $Nu_{(247\text{-}419)}$ signal before and after binding to s2m RNA. The signals stemming from disordered regions remain NMR observable upon binding. **b** 1D DOSY spectra acquired for multiple gradient strengths on $Nu_{(247\text{-}419)}$ condensate sample formed with s2m RNA. The last spectrum acquired with 45 G/cm gradient strength shown in black stems from the protein in condensed phase. Note that the IDR peaks have very similar linewidth in both condensed and dilute phases illustrating that longer disordered regions remain flexible even in the droplets.

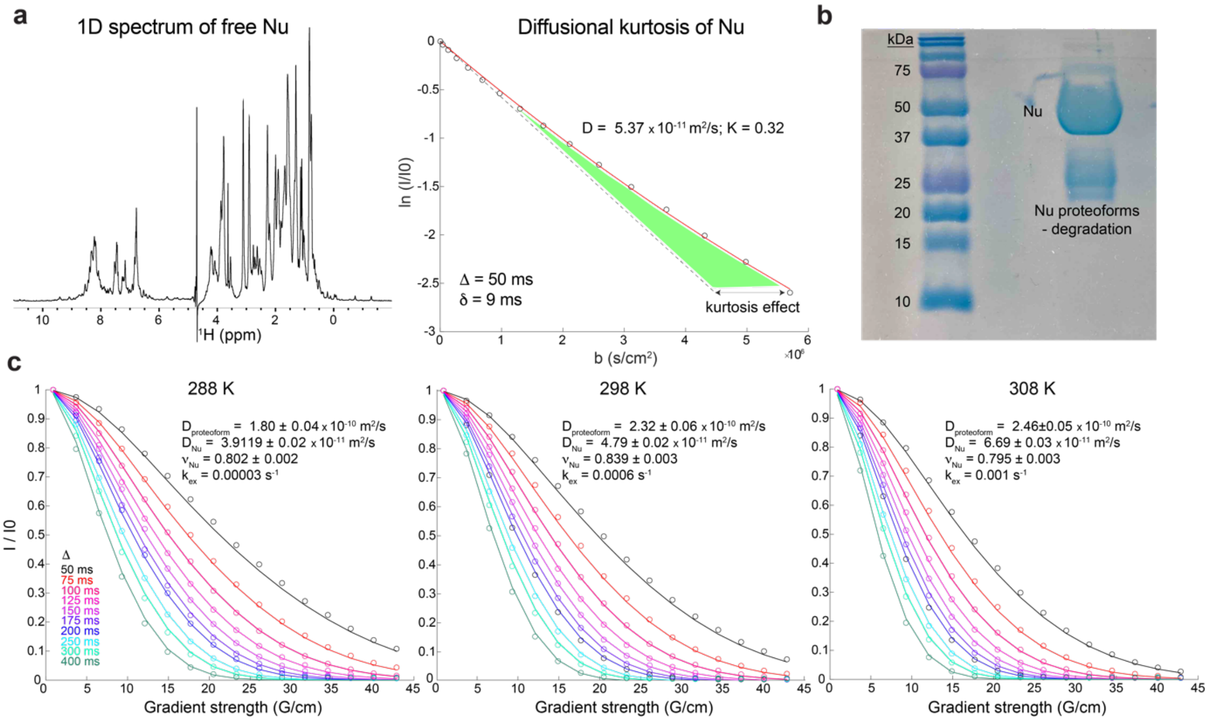


**Supplementary Fig. 6: LLPS REDIFINE can also detect protein degradation. a** $^{1}$H spectrum of a free Nucleocapsid protein after a couple of weeks at the room temperature showing at first sight no signs of degradation. However, diffusional kurtosis is observed suggesting the presence of more than one species in solution. **b** On the SDS-PAGE gel it is clear that sample contains also another Nucleocapsid proteoform of around 25 kDa in molecular weight. **c** LLPS REDIFINE data acquired on this degraded Nucleocapsid sample. Two species attributed to free Nucleocapsid and a smaller proteoform are identified. More important, no exchange between two species is detected which is consistent with the existence of two independent proteins in solution. This is confirmed by recording the experiments at 288, 298 and 308 K. All fitting results show constant populations and no chemical exchange between the species.
